# TEDLH: Domain HMMs for sensitive detection of remote homologues

**DOI:** 10.64898/2026.01.07.697892

**Authors:** Claudia Alvarez Carreño, Anton S. Petrov, Vaishali P. Waman, Ian Sillitoe, Christine Orengo

**Affiliations:** Department of Structural and Molecular Biology, University College London, London, United Kingdom; NASA Center for the Origin of Life, Georgia Institute of Technology, Atlanta, GA 30332-0400, USA; School of Chemistry and Biochemistry, Georgia Institute of Technology, 901 Atlantic Dr, Atlanta, GA 30332, USA

## Abstract

**Motivation:** The Encyclopedia of Domains (TED) provides domain annotations for proteins in the AlphaFold Protein Structure Database (AFDB) using a consensus of three state-of-the-art structure-based methods. We used these TED domain annotations to construct profile Hidden Markov models (HMMs), collectively forming the TED Library of HMMs (TEDLH). TEDLH enables sensitive sequence and profile searches, supporting systematic exploration of protein domain families and their evolutionary relationships.

**Results:** TEDLH links domain HMMs to experimentally determined CATH-PDB structures through direct (primary) and transitive (secondary and tertiary) relationships. Fewer than half of TEDLH HMMs are directly linked to a CATH-PDB domain; the remaining models are connected through transitive relationships. These transitive links extend coverage into more divergent regions of sequence space and better represent CATH superfamily diversity.

HMM–HMM comparisons within CATH superfamily 3.30.70.100 illustrate how transitive relationships expand sequence coverage in TEDLH. In this superfamily, 4,813 TEDLH HMMs are connected to 212 CATH-PDB representatives. Primary, secondary, and tertiary relationships progressively capture more divergent sequences (pairwise sequence identity <20%) that retain structural similarity (TM-score >0.6) and a conserved two-layer α/β sandwich core fold.

All-against-all HMM–HMM comparisons across TEDLH also reveal sequence similarities across the CATH hierarchy (cross-hits). At low query coverage (<50%), cross-hits are more frequent between CATH classes, whereas at higher coverage thresholds (>70%) they predominantly occur between superfamilies. These cross-hits are not driven by superfamily size or sequence diversity and can provide guidance for CATH curation. As an example, analysis of cross-hits between superfamilies 2.170.130.30 and 3.10.20.30 reveals evolutionary relationships between these groups.

**Availability and Implementation:** TEDLH is compatible with HH-suite3 and is available from FigShare https://doi.org/10.6084/m9.figshare.28531754 for local use.

## Background

Protein domains are conserved units of function and evolution. For the past thirty years, domain annotation has followed two complementary paths. Domains identified in experimentally determined structures from the Protein Data Bank (Berman, et al., 2000) have been systematically classified and mapped onto protein sequences by resources such as CATH (Orengo, et al., 1997), SCOP (Murzin, et al., 1995), and ECOD (Cheng, et al., 2014). Parallel approaches based on alignments of homologous sequences have enabled the identification of domains in protein sequences through patterns of sequence conservation (e.g. Pfam (Sonnhammer, et al., 1997), SMART (Schultz, et al., 1998), and TIGRFAMS (Haft, et al., 2003)). Together with structure-based classifications, these efforts laid the foundation for profile-based models, including Gene3D (Buchan, et al., 2002) and SUPERFAMILY (Gough and Chothia, 2002), which extend domain annotation to millions of uncharacterized sequences while capturing both evolutionary and structural information. More recently, highly accurate protein structure prediction methods have provided access to predicted structures at an unprecedented scale. The availability of this expanded structural data creates new opportunities, as well as new challenges, for structure-informed domain annotation and classification.

The AlphaFold Database (AFDB) (Varadi, et al., 2022) transformed access to protein structural data by supplying predicted three-dimensional models for more than 214 million UniProt (UniProt_Consortium, 2019) sequences. Since its release, several resources have been developed to provide annotation and to integrate AFDB with other bioinformatics databases and resources. The Encyclopedia of Domains (TED) (Lau, et al., 2024) catalogues 365 million putative domains in proteins from AFDB (version 4), of which more than 251 million are mapped to CATH (Orengo, et al., 1997). CATH (Orengo, et al., 1997) is one of the most widely used protein domain classification systems, organizing domains hierarchically into Class, Architecture, Topology, and Homologous superfamily to capture shared structural features and evolutionary relationships. Domain assignments in TED are derived from the consensus of three state-of-the-art structure-based methods: Merizo (Lau, et al., 2023), Chainsaw (Wells, et al., 2024), and UniDoc (Zhu, et al., 2023). TED greatly expands both the number and the sequence diversity of domains that can be assigned to CATH with high confidence.

Here, we present TEDLH (TED Library of Hidden Markov Models). Data for TEDLH were derived from non-redundant high-quality TED domains, as determined by the Qscore, which integrates multiple indicators of assignment reliability. This strategy propagates the structure-based domains annotated in TED to the sequence level. By retrieving sequences directly from TED and using its pre-defined domain boundaries, we mitigate the risk of homologous over-extension (Gonzalez and Pearson, 2010) that can occur in iterative sequence searches (e.g., PSI-BLAST (Altschul, et al., 1997) or jackhammer (Johnson, et al., 2010)). Moreover, because TED annotates the AFDB (version 4), it provides a broad and diverse set of domain sequences. As a result, sequence diversity can be sourced directly from TED to build multiple sequence alignments (MSAs) rather than from general protein sequence databases. These features give TED-derived MSAs two major advantages: precise domain boundary definitions and high sequence diversity.

Here we use similarity to intermediate sequences to map experimentally determined structures in the PDB (Berman, et al., 2000) classified by CATH (CATH-PDB) to a library of HMMs built from TED sequences (TEDLH HMMs). Intermediate sequence searches, have long been employed to detect remote homologous relationships because these methods substantially increase the sensitivity of sequence comparisons (Li, et al., 2000; Park, et al., 1997; Salamov, et al., 1999). HMM identifiers in TEDLH encode the associated CATH-PDB domain and superfamily, as well as whether the connection is direct (primary) or through an intermediate HMM (secondary or tertiary).

To demonstrate the functionality of the TEDLH library, we selected two examples from the all-against-all scanning of HMMs in TEDLH. The first example illustrates sequence relationships within superfamily 3.30.70.100. We also show examples of ancestral relationships across CATH superfamilies 2.170.130.30 and 3.10.20.30. TEDLH facilitates efficient exploration of protein domain diversity at the sequence level. TEDLH is publicly available via figshare.

## Methods

### Domain selection and initial library preparation

Sequences of proteins with non-redundant domains in the TED database (Lau, et al., 2024) were retrieved from UniProt and trimmed according to the domain boundaries defined in TED. Each domain sequence was labelled with its corresponding CATH superfamily assignment in TED and subsequently filtered by Qscore. The Qscore assesses the reliability of a TED domain assignment by integrating structural coverage, domain assignment consensus, compactness, and pLDDT. It ranges from 0 (no confidence) to 100 (high confidence). Qscores above 75 are considered high confidence. High-confidence TED domains were grouped according to their structure-based CATH superfamily assignments provided by TED. TED sequences within each CATH superfamily group were then clustered by sequence similarity with mmseqs (Hauser, et al., 2016) using a maximum *E*-value threshold of 1×10^−3^ and a bi-directional coverage threshold of 80. This strategy takes advantage of the high-quality domain boundaries and the inclusion of remote relatives defined by TED. Clustering with mmseqs yielded 945,404 singletons and 889,249 clusters with two or more sequences.

Because the full potential of an HMM comes from the variation captured across multiple homologous sequences, singletons were excluded, and the remaining clusters were each aligned with MAFFT (Katoh and Standley, 2013). For clusters with more than 300 members, redundancy was reduced by selecting representative sequences at a maximum of 70% pairwise identity using mmseqs clustering.

Multiple sequence alignments (MSAs) were converted into individual HMMs using hhbuild from HH-suite3 (Steinegger, et al., 2019). To make all-against-all HMM–HMM comparisons computationally tractable, the 889,249 HMMs were partitioned into six tailored libraries following the HH-suite3 user guide, omitting secondary structure scoring as recommended. These libraries comprised: (i) CATH class 1 (176,678 HMMs); (ii) CATH class 2 (307,438 HMMs); (iii) class 3, architecture 3.30 (2-layer sandwich; 152,952 HMMs); (iv) class 3, architectures 3.40, 3.50 and 3.55 (3-layer sandwiches; 138,851 HMMs); (v) superfamily 3.40.50.300 (P-loop NTP hydrolases; 23,389 HMMs); and (vi) remaining class 3 architectures (3.10, 3.15, 3.20, 3.60–3.100; 89,941 HMMs). All 889,249 HMMs were queried against each library using hhblits. All-against-all HMM–HMM comparison results were analysed to identify transitive relationships between HMMs and to detect cross-hits between distinct CATH superfamilies.

### TEDLH Library curation

#### Recovering TED-HMMs related to CATH

To ensure that the final library contained only TED-HMMs with unambiguous links to experimentally determined CATH-PDB domains, we performed a cross-referencing step. Non-redundant CATH-PDB domain sequences (CATH v4_4_0) were scanned against all TED-HMMs using HHblits (Steinegger, et al., 2019). A TED-HMM was considered to have a primary relationship to a CATH-PDB domain only if two criteria were met: (i) the HMM–sequence match exceeded predefined thresholds (E-value ≤ 1×10^−1^, query coverage ≥ 70%, template coverage ≥ 70%), and (ii) the TED domains used to build the HMM were all assigned by TED to the same CATH superfamily as the matched CATH-PDB domain. We adopted a relaxed E-value threshold because the agreement between independent structure-based TED assignments and CATH-PDB classifications provides additional support for homology. For each primary TED-HMM, the best-matching CATH-PDB domain was retained and encoded in the HMM header together with the corresponding CATH superfamily (Figure 1b).

**Figure 1.**
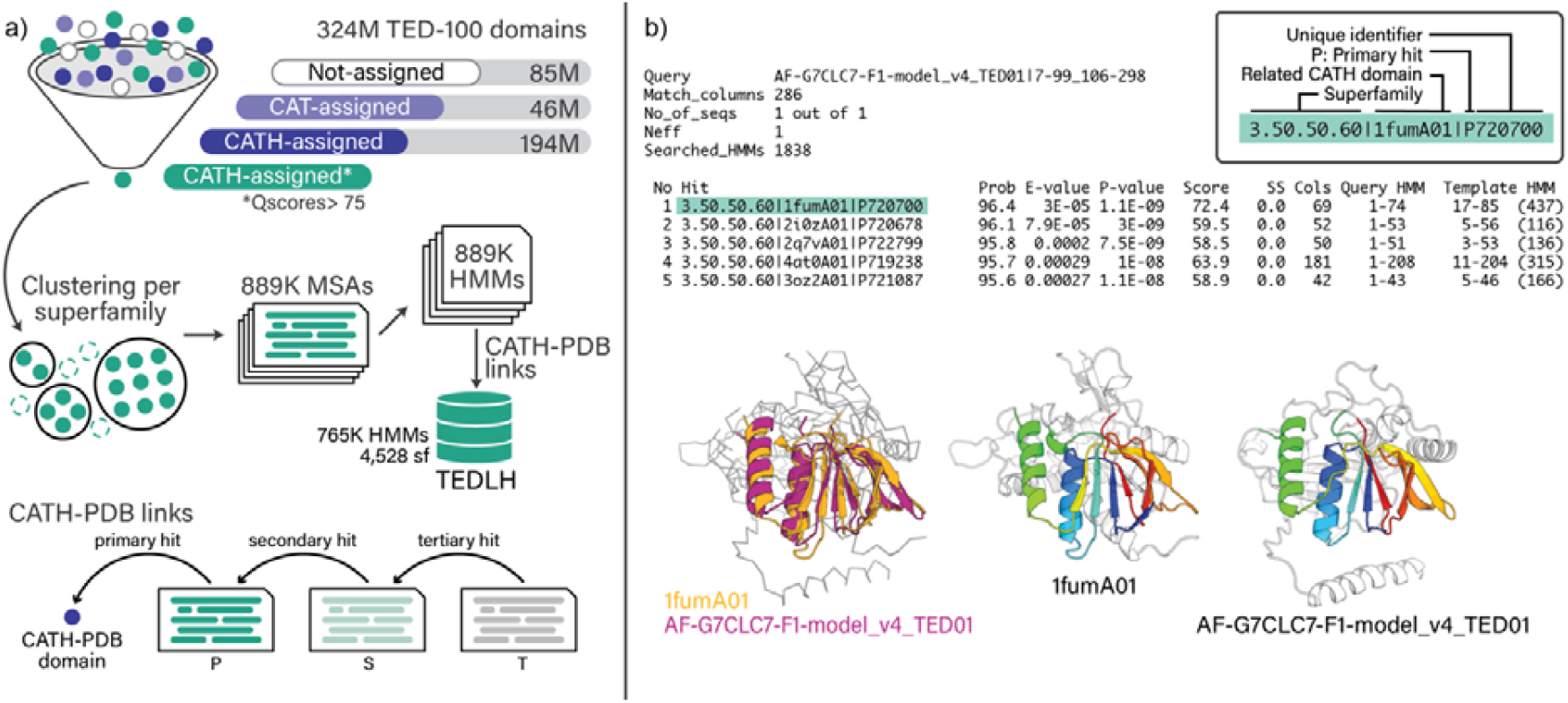
TED Library of HMMs. (a) Construction of TEDLH from TED domain annotations. TED domain sequences were grouped by CATH superfamily and clustered by sequence similarity using MMseqs. Sequences within each cluster were aligned to generate multiple sequence alignments (MSAs), which were subsequently converted into profile hidden Markov models (HMMs). TEDLH comprises 765,248 TED-HMMs with primary, secondary, or tertiary relationships to CATH-PDB domains. (b) Example output from an HHblits search against TEDLH. Regions of structural similarity between the query and the corresponding CATH-PDB template are highlighted in colour. Inset: information encoded in the HMM header, including the associated CATH superfamily (sf).

To identify TED-HMMs related to CATH-PDB beyond primary matches, we analysed all-against-all HMM–HMM comparisons within TED-HMMs. Secondary relations were defined as TED-HMMs linked to a primary HMM that matches a CATH-PDB domain, and tertiary relations were defined as TED-HMMs linked only to secondary HMMs. The same thresholds used for primary hits (E-value ≤ 1×10^−1^, query and template coverage ≥70%) were applied for secondary and tertiary relations. Additionally, all sequences in the TED-HMM MSA were required to share the same CATH classification as the CATH-PDB domain to which they are ultimately linked. TED-HMMs with primary, secondary, or tertiary relations to CATH-PDB were compiled into the final TED HMM Library (TEDHL). Using this strategy, we recovered 765,248 TED-HMMs that could be associated with a CATH-PDB domain through transitive homologous relationships. Thus, primary, secondary and tertiary relationships reflect both sequence similarity and concordant structure-based domain classification.

The final library (TEDLH) was built with the HH-suite3 using these 765,248 TED-HMMs, divided into three sets of i) class 1 (168,863 HMMs spanning 1,559 CATH superfamilies); ii) class 2 (230,774 HMMs spanning 1,043 CATH superfamilies); and iii) class 3 (365,611 HMMs spanning 1,926 CATH superfamilies). Overall, TEDHL HMMs map to 51,180 CATH-PDB sequences from 4,528 CATH superfamilies, representing approximately 80% of superfamilies in CATH v4.4.

### Sequence and structure analyses

Cluster map visualizations of relationships within TEDLH HMMs were done with CLANS (Frickey and Lupas, 2004) using E-values from precomputed all-against-all scans.

Pairwise comparisons at the sequence level (HHM-HHM) were performed with hhalign from the HHsuite (Steinegger, et al., 2019), and at the structure level with TM-align (Zhang and Skolnick, 2005). Structure-derived MSAs were calculated with MATRAS (Kawabata, 2003). We used CATH-SSAP for superpositions of multiple structures (Orengo and Taylor, 1996). Topology diagrams were downloaded from ProteoVision (Penev, et al., 2021).

## Results

### Contribution of primary and transitive links to TEDLH coverage of CATH superfamilies

The TEDLH library comprises HMMs derived from TED domain sequences that can be associated with CATH-PDB domains either directly or through intermediate HMMs. We first identified TEDLH HMMs exhibiting direct sequence similarity with CATH-PDB domains (primary relations). TEDLH HMMs that display significant similarity to these primary HMMs were then collected as secondary relations. Finally, we retrieved HMMs related to the set of secondary relations (tertiary relations). This series of transitive connections enable the detection of remote homologs (Li, et al., 2000; Park, et al., 1997; Salamov, et al., 1999). The HMM identifier indicates the level of the transitive relationship to a CATH-PDB domain and the corresponding CATH superfamily (Figure 1b).

For each CATH superfamily, we assessed how well TEDLH models capture the sequence space associated with experimentally determined structures. Primary relationships are detected through direct searches of CATH-PDB domains against the TEDLH HMM library, whereas secondary and tertiary relations encompass more divergent homologues, thus, extending coverage into more remote regions of the sequence space. We quantified the proportion of TEDLH HMMs that are directly linked to a CATH-PDB domain (primary relationships) across CATH structural classes. In CATH class 1, 42.1% of TEDLH HMMs are associated with a primary relationship to a CATH-PDB domain. In class 2, primary relationships account for 28.4% of TEDLH HMMs. In class 3, 47.4% of TEDLH HMMs are directly linked to a CATH-PDB domain. The remaining coverage is contributed by secondary and tertiary relationships. Secondary and tertiary relations broaden the mapping of sequence space so that the diversity of domains in CATH can be explored by transitive relationships.

### Cross-hits across CATH hierarchy levels

All-against-all HMM–HMM comparisons in TEDLH also recover sequence similarity across-superfamilies (cross-hits) (Figure 2c). These cross-hits can occur between superfamilies within a different CATH Class, Architecture, Topology or Homology. At an E-value threshold of 1×10^−3^, the number of superfamilies that retrieve cross-hits varies depending on the minimum query coverage threshold. Between 20% and 50% query coverage thresholds, cross-hits between classes are more frequent. At higher query coverage thresholds (above 70% query coverage), most cross-hits occur between superfamilies and may indicate previously undetectable relationships; detection of these sequence similarities can offer valuable guidance for the manual curation of CATH. The top 25 superfamilies with the most cross-hits are shown in Figure 2d and Table 1. The number of cross-hits does not appear to be driven by superfamily size or sequence diversity, as no clear pattern is seen with the number of HMMs or the number of S95/S35 clusters in CATH (Table 1).

**Table 1.** Diversity of the top 25 CATH-PDB superfamilies with the most cross-hits at 80% coverage.

| **Superfamily (name)** | **HMMs in TEDLH** | **Domains in CATH** | **S95 in CATH** | **S35 in CATH** |
| --- | --- | --- | --- | --- |
| 2.60.40.10<br>(Immunoglobulins) | 42,591 | 31,905 | 4,547 | 873 |
| 2.60.40.1930<br>(Macroglobulin (MG2)<br>domain) | 4,434 | 181 | 19 | 13 |
| 3.40.50.2300<br>(Response regulator) | 11,110 | 2,543 | 616 | 402 |
| 1.10.10.10 (Winged<br>helix-like DNA-binding<br>domain<br>superfamily/Winged<br>helix DNA-binding<br>domain) | 29,985 | 3,444 | 768 | 524 |
| 3.30.70.260 (ACT<br>domain) | 2,320 | 335 | 87 | 50 |
| 3.40.50.720 (NAD(P)-<br>binding Rossmann-like<br>Domain) | 9,283 | 11,728 | 1,430 | 647 |
| 3.30.70.100 | 4,813 | 723 | 168 | 101 |
| 2.60.40.1220 | 2,155 | 64 | 15 | 10 |
| 2.60.40.2550 | 963 | 2 | 1 | 1 |
| 2.40.50.140 (Nucleic<br>acid-binding proteins) | 9,368 | 2,879 | 415 | 227 |
| 3.40.50.300 (P-loop<br>containing nucleotide<br>triphosphate<br>hydrolases) | 22,877 | 9,233 | 1,310 | 561 |
| 2.60.40.1080 | 2,099 | 91 | 31 | 13 |
| 2.30.30.140 | 1,671 | 641 | 130 | 77 |
| 2.60.40.650 | 538 | 43 | 6 | 4 |
| 2.60.40.1120<br>(Carboxypeptidase-like,<br>regulatory domain) | 6,693 | 46 | 15 | 10 |
| 3.40.50.2000 (Glycogen | 5,451 | 1,605 | 190 | 103 |
| Phosphorylase B) |  |  |  |  |
| 2.60.120.260<br>(Galactose-binding<br>domain-like) | 9,795 | 1,427 | 144 | 236 |
| 2.60.40.3880 | 346 | 40 | 2 | 2 |
| 3.20.20.70 (Aldolase<br>class I) | 2,955 | 6,031 | 603 | 233 |
| 2.60.40.2950 | 126 | 2 | 1 | 1 |
| 3.30.70.120 | 503 | 451 | 56 | 23 |
| 3.30.420.40 (ATPase,<br>nucleotide binding<br>domain) | 2,584 | 2,589 | 316 | 166 |
| 2.60.40.740 | 886 | 75 | 25 | 17 |
| 2.60.40.2310 | 486 | 7 | 3 | 2 |
| 2.30.30.30 | 695 | 619 | 71 | 32 |

**Figure 2.**
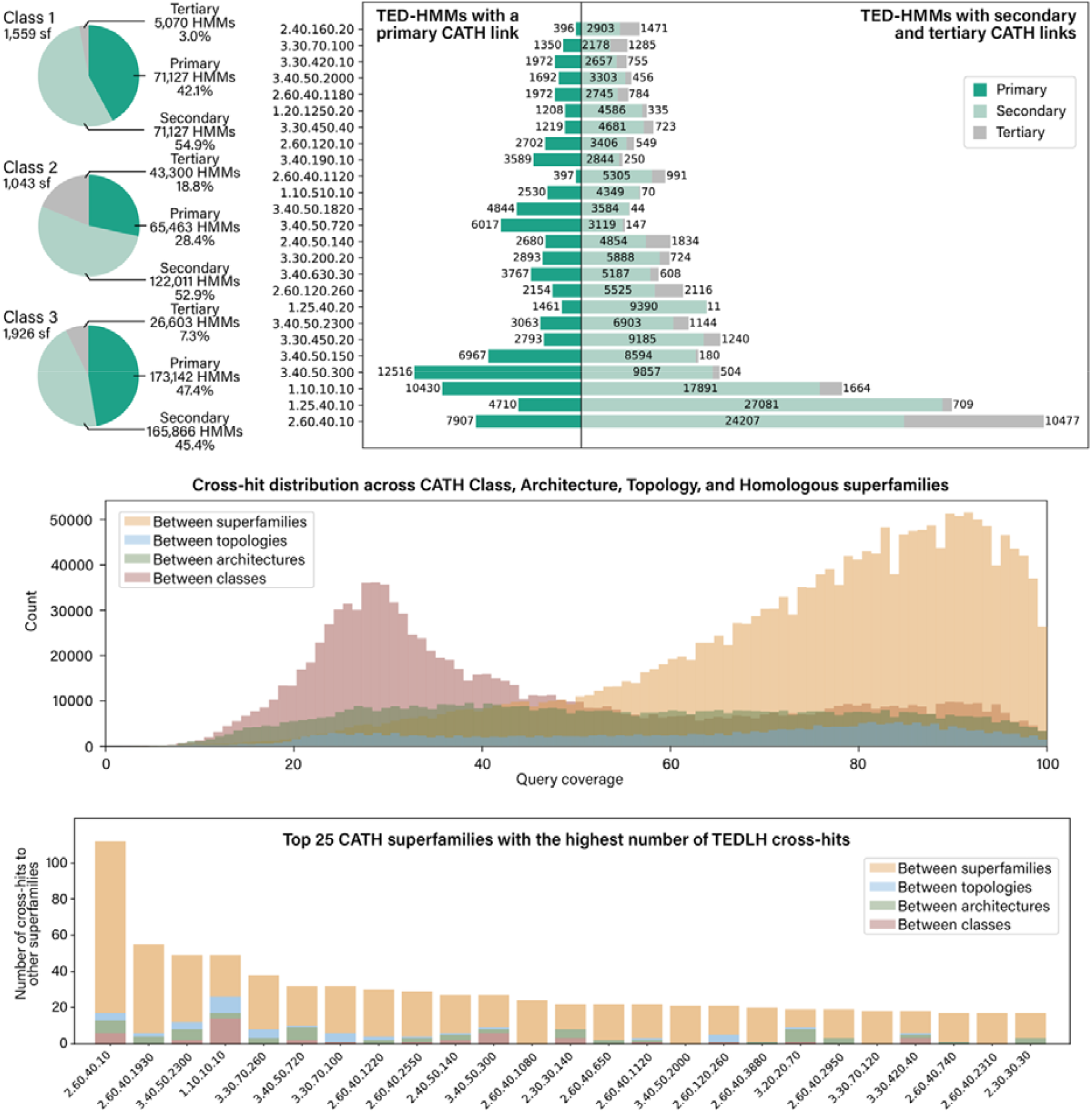
TED-HMM relationships to CATH-S100 and cross-hits across TEDHL superfamilies. (a) Proportions of TED-HMMs exhibiting primary, secondary, or tertiary connections to CATH-PDB domains, shown separately for class 1, class 2, and class 3. (b) Distribution of primary, secondary, and tertiary TED-HMM relationships to CATH-S100 domains for the 25 most represented superfamilies in TEDLH. (c) Histogram of TEDLH cross-hits across different levels of the CATH hierarchy at varying query coverage thresholds. (d) The 25 CATH superfamilies with the highest number of TEDLH cross-hits at a 70% bi-directional coverage threshold.

Cross-superfamily sequence similarities reveal remote evolutionary relationships. The superfamily with that cross-hits the largest number of other superfamilies in TEDLH at a 70% bi-directional coverage threshold is 2.60.40.10, Immunoglobulins (Figure 2d). HMMs of superfamily 2.60.40.10 cross hit 112 other superfamilies, of these, 95 cross-hits are between CATH Homology levels; 4 are between CATH Topologies levels; 7 are between CATH Architectures; and 6 between CATH Classes.

### Homology relationships within superfamily 3.30.70.100

To illustrate the ability of TED-HMMs to extend sequence coverage, we analysed 4,813 models within superfamily 3.30.70.100. These HMMs are linked, directly or via transitive connections, to 212 non-redundant CATH-PDB domains. A cluster map representation was generated from precomputed pairwise HMM–HMM comparisons among HMMs and between CATH-PDB sequences and HMMs (Figure 2a). By design, at a relaxed E-value threshold of 1×10^−1^, all TED-HMMs are linked to at least one CATH-PDB representative (Figure 3a).

**Figure 3.**
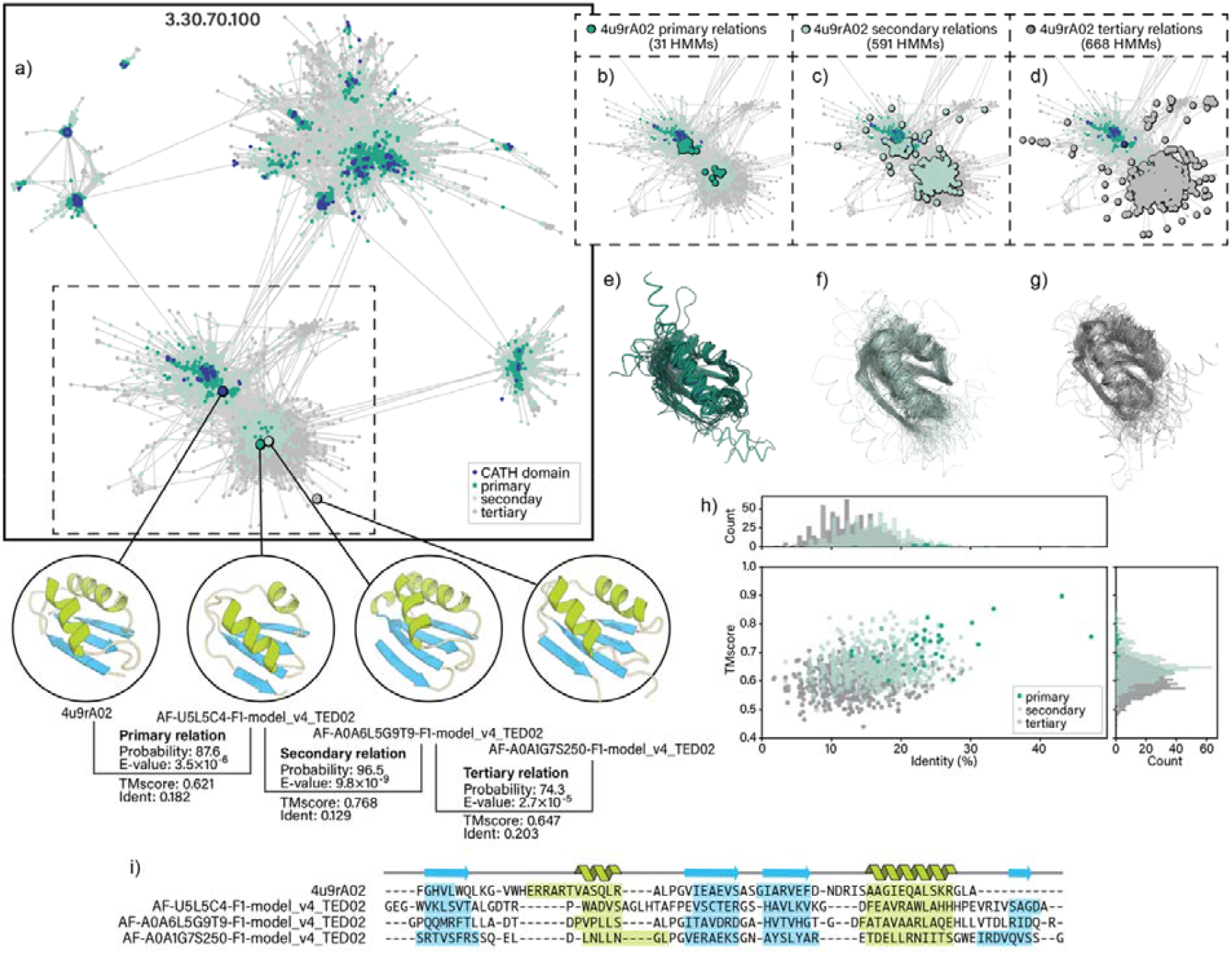
Represenation of primary secondary and tertiary relationships between HMMs and CATH S100 domains of superfamily 3.30.70.100. (a) Cluster map of HMM-HMM similarities for superfamily 3.30.70.100 showing connections at an *E*-value threshold of 1×10^−1^. TED domains AF-U5L5C4-F1-model_v4_TED02, AF-A0A6L5G9T9-F1-model_v4_TED02, AF-A0A1G7S250-F1-model_v4_TED02. (b–d) Same map, highlighting primary (b), secondary (c), and tertiary (d) connections to CATH domain 4u9rA02. (e) Structural superposition of CATH domain 4u9rA02 with one representative from each of its 31 primary HMMs. (f) Structural superposition of CATH domain 4u9rA02 with representatives from its 591 secondary HMMs. (g) Structural superposition of CATH domain 4u9rA02 with representatives from its 668 tertiary HMMs. (h) Relationship between structural similarity (TM-score) and sequence identity for 4u9rA02 compared with representatives of its primary, secondary, and tertiary HMMs. (i) Structure-informed multiple sequence alignment of domains 4u9rA02, AF-U5L5C4-F1-model_v4_TED02, AF-A0A6L5G9T9-F1-model_v4_TED02, AF-A0A1G7S250-F1-model_v4_TED02 with secondary structural elements.

We examined the set of TEDLH relationships associated with the CATH-PDB domain 4u9rA02 (superfamily 3.30.70.100), a metal-binding domain from the *Cupriavidus metallidurans* P1B-4-ATPase CzcP protein (Smith, et al., 2015). In total, 31 primary, 591 secondary, and 668 tertiary relations were identified (Figure 3a-d). A superposition of 4u9rA02 to representatives of primary, secondary, and tertiary connections shows high structure similarity (TMscores > 0.6) despite low pairwise sequence identities (Sequence identity < 20%). Primary, secondary, and tertiary HMMs progressively sample more distant homologues, capturing sequences with lower pairwise identity and greater structural variation. Despite this divergence, structural superpositions across multiple representatives reveal a conserved core fold, consisting of a β-α-β-β-α-β arrangement that forms a two-layer α/β sandwich. The analysis of superfamily 3.30.70.100 illustrates the ability of TED-HMMs to detect remote homology beyond regions covered by experimentally determined structures.

### Insights into Protein Fold Divergence

#### Evolutionary fold switching

To investigate potential evolutionary relationships between distinct folds, we examined cross-superfamily HMM–HMM similarities. Comparisons between CATH superfamilies 2.170.130.30 and 3.10.20.30 reveal significant sequence similarity (HHalign Probability 90%, E-value 4.9×10^−7^), suggesting a shared evolutionary origin (Figure 4). Although domains in these superfamilies show detectable sequence similarity, their differences emerge at a higher level of structural organization.

**Figure 4.**
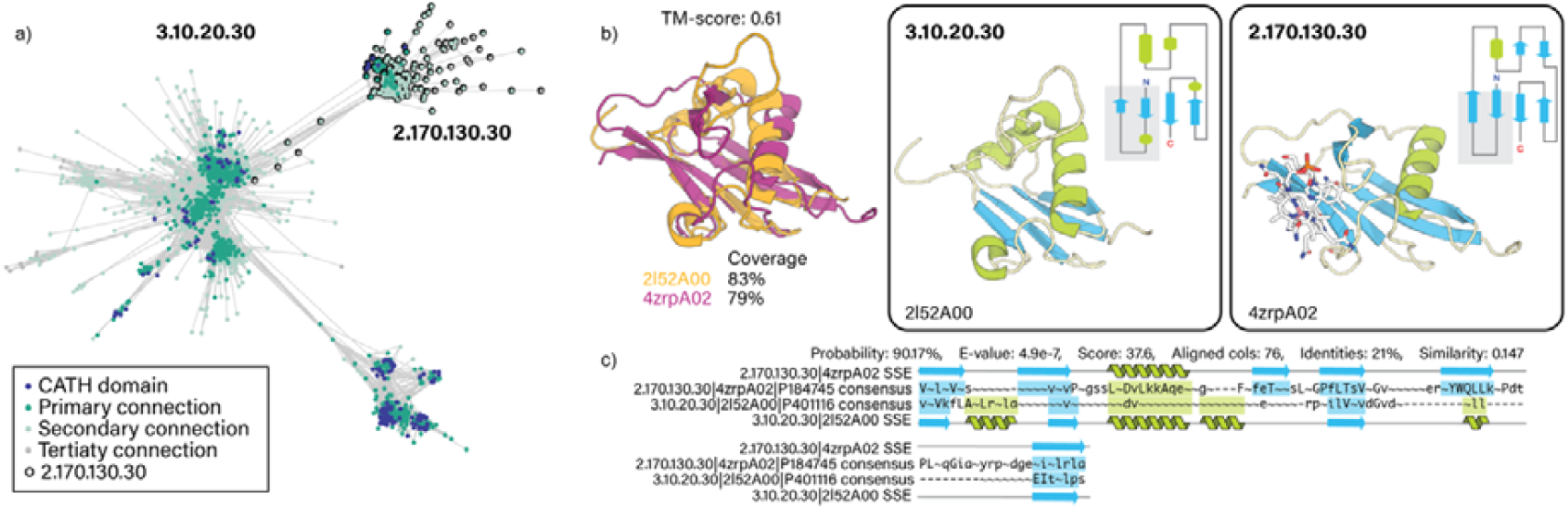
Evolutionary fold switching. (a) Cluster map of HMM-HMM similarities between superfamilies 2.170.130.30 and 3.10.20.30 showing connections at an E-value threshold of 1×10^−3^. (b) Structure superposition of 2l52A00 (representative of superfamily 2.170.130.30) and 4zrpA02 (representative of superfamily 3.10.20.30). Insets: three-dimensional structure and topology diagrams of domains 2l52A00 and 4zrpA02 coloured by secondary structural element. (c) HMM-HMM alignment.

Pairwise structural comparisons between 2l52A00 (representative of superfamily 3.10.20.30) and 4zrpA02 (representative of superfamily 2.170.130.30) show overall structural similarity (TM-score 0.61). The observed differences arise mainly from changes in the composition and arrangement of secondary-structure elements (SSEs). In domain 2l52A00, the SSEs are organised as β(2)–[α]– β(2), whereas in domain 4zrpA02 they adopt a β(2)–[αββ]–β (2) arrangement. In both cases, the N- and C-terminal β-hairpins form a comparable β-sheet that superimposes well, while the regions highlighted in brackets represent the variable structural segment that distinguishes the two domains (Figure 4b-c). This pattern is consistent with a possible interconversion between disorder and secondary structural elements during evolution. CATH superfamily 2.170.130.30 includes several cobalamin-binding proteins, such as transcobalamin-1 (4kkiA02), transcobalamin-2 (2bb6A02, 4zrpA02, 5nsaA00), and intrinsic factor (2pmvA02). These domains share a characteristic β-hairpin, which is absent from those in 3.10.20.30. The β-hairpin provides a surface involved in the coordination of cobalamin (Figure 4).

The results of pairwise comparisons of HMMs in TEDLH suggests that these superfamilies, currently placed in different classes, could be considered for merging in CATH. The class distinction is based on the proportion of residues in helix versus strand conformations: 2.170.130.30 is assigned to class 2 (mainly β) due to a higher proportion of β-strand residues and a lower proportion of α-helices compared to 3.10.20.30.

## Discussion

The identification of domains at the structure level in TED brings in a wealth of annotation for domains that would have been very difficult to identify by sequence alone (Lau, et al., 2024). Domains in TED are identified based on three complementary domain-chopping methods (Merizo (Lau, et al., 2023), Chainsaw (Wells, et al., 2024) and UniDoc (Zhu, et al., 2023)), providing precise and reliable domain boundaries, which we used to generate MSAs that retain structural information and serve as a robust source of evolutionary data. Using these annotations, we generated high-quality sequence HMMs, creating TEDLH, a library derived from MSAs of sequence clusters covering non-redundant TED domains assigned to CATH.

TEDLH currently contains high-confidence domains mapped to CATH and linked, directly or via transitive relationships, to a CATH-PDB structure. Each TEDLH HMM can be traced to its closest CATH-PDB representative, facilitating evolutionary and functional interpretation and downstream annotation. By including remote homologues through transitive links, TEDLH captures structural similarity (TM-score > 0.50) even when sequence identity is low (<20%). In contrast to purely structure-based comparisons and embeddings-based comparisons, which require additional evidence to infer evolutionary relationships, sequence matches against TEDLH are based on conserved amino-acid positions and insertion/deletion patterns, providing directly interpretable signals of shared ancestry.

The patterns of partial cross-hits (query coverage < 50%) observed for a large proportion of superfamilies represented in TEDLH is consistent with previous observations that protein folds often share localized regions of similarity (Alva, et al., 2015; Ferruz, et al., 2020; Friedberg and Godzik, 2005; Kolodny, et al., 2021). These local regions of similarity likely reflect remnants of shared evolutionary history between structurally distinct folds (Alva, et al., 2015; Fetrow and Godzik, 1998; Kolodny, et al., 2021; Lupas, et al., 2001). Global cross-hits (query and template coverage ≥ 70%) likely reflect remote homology relationships that become detectable with the expanded sequence coverage offered by TEDLH profiles.

TEDLH models are searchable with HHblits and HHsearch from HH-suite3 (Steinegger, et al., 2019), allowing both single-sequence queries and HMM–HMM comparisons. TEDLH complements established HMM-based resources for sequence similarity detection at the domain level such as the Pfam protein family database (Paysan-Lafosse, et al., 2025) (harmonized with the ECOD structural classification (Cheng, et al., 2014)) and Gene3D (based on the CATH classification of protein structures). TEDLH extends access to AFDB-derived structural diversity at the sequence level and provides a valuable resource for investigating remote homology and informing the ongoing curation of domain classifications. TEDLH will be expanded in the future to incorporate additional superfamilies not yet classified in structural databases, further increasing its coverage and utility for detecting remote homology.

## Data availability

TEDLH is publicly available and downloadable from figshare https://doi.org/10.6084/m9.figshare.28531754.

## Acknowledgements

CAC is funded by the Royal Society Newton International Fellowship [NIF\R1\231037]. ASP acknowledges funding from NASA Grant 80NSSC24K0344. VPW is funded by Wellcome Trust [221327/Z/20/Z]. IS is funded by BBSRC [BB/W018802/1].

